# Multi-modal meta-analysis of 1494 hepatocellular carcinoma samples reveals vast impacts of consensus driver genes on phenotypes

**DOI:** 10.1101/166090

**Authors:** Kumardeep Chaudhary, Olivier B Poirion, Liangqun Lu, Sijia Huang, Travers Ching, Lana X Garmire

**Affiliations:** Epidemiology Program, University of Hawaii Cancer Center, Honolulu, HI 96813, USA; Molecular Biosciences and Bioengineering Graduate Program, University of Hawaii at Manoa, Honolulu, HI 96822, USA

## Abstract

Although driver genes in hepatocellular carcinoma (HCC) have been investigated in various previous genetic studies, prevalence of key driver genes among heterogeneous populations is unknown. Moreover, the phenotypic associations of these driver genes are poorly understood. This report aims to reveal the phenotypic impacts of a group of consensus driver genes in HCC. We used MutSigCV and OncodriveFM modules implemented in the IntOGen pipeline to identify consensus driver genes across six HCC cohorts comprising 1,494 samples in total. To access their global impacts, we used TCGA mutations and copy number variations to predict the transcriptomics data, under generalized linear models. We further investigated the associations of the consensus driver genes to patient survival, age, gender, race and risk factors. We identify 10 consensus driver genes across six HCC cohorts in total. Integrative analysis of driver mutations, copy number variations and transcriptomic data reveals that these consensus driver mutations and their copy number variations are associated with majority (62.5%) of the mRNA transcriptome, but only a small fraction (8.9%) of miRNAs. Genes associated with *TP53, CTNNB1*, and *ARID1A* mutations contribute to the tripod of most densely connected pathway clusters. These driver genes are significantly associated with patients’ overall survival. Some driver genes are significantly linked to HCC gender (*CTNNB1, ALB, TP53* and *AXIN1*), race (*TP53* and *CDKN2A*), and age (*RB1*) disparities. This study prioritizes a group of consensus drivers in HCC, which collectively show vast impacts on the phenotypes. These driver genes may warrant as valuable therapeutic targets of HCC.

## Introduction

Liver cancer is the leading cause of cancer deaths worldwide, with more than 700,000 incidences and deaths in recent years (1). Globally, this cancer is ranked second for cancer-related mortality among men (2). In the US, it is one of the few cancers with increased rate of ∼3% per year, for both incidence and mortality (3). Hepatocellular carcinoma (HCC) is the prominent histological type of liver cancer and accounts for approximately 75%-90% of all the liver cancer cases (4). The incidence rates of HCC vary by factors such as race, gender, age as well as demographic regions. East Asians are twice likely to develop liver cancer compared to Caucasian or African American populations (5). Additionally, males have 2 to 4 times higher incidence rates than females. The incidence rates peak around 60-65 years for males and 65-70 for females (6,7). Various other risk factors for the HCC development have been well-determined, such as cirrhosis, hepatitis B (HBV) infection, hepatitis C (HCV) infection, alcohol abuse, obesity and environmental toxic intake (8). While HBV infection is the major risk for HCC cases in East Asian countries, HCV and alcohol abuse are the leading causes of HCC in North America and Europe (9).

The initiation and advancement of cancer are thought to occur after continuous accumulations of somatic genomic alterations, followed by the widespread manifestation of gene products (10 – 13). Using the whole genome sequencing (WGS) or whole exome-sequencing (WES) technology, many studies have aimed to determine candidate driver gene mutations in HCC, the type of mutations that confer a selective growth advantage to the cell (14 – 20). *TP53* and *CTNNB1* are reported as the two most frequently mutated genes in HCC (21). Other putative driver genes include those related to genome stability, such as *ARID1A, ARID2*, and *MLL1-4* (15,17,22 – 24), *RB1* in cell cycle pathway (16), *AXIN1* in Wnt signaling pathway (25), *NFE2L2* in oxidative stress (22), and *TSC1/TSC2* in MAPK signaling pathway (16,22). A recent analysis of hepatocellular carcinoma from The Cancer Genome Atlas (TCGA) reported the significant mutation of *LZTR1* (encoding an adaptor of CUL3-containing E3 ligase complexes) and *EEF1A1* (encoding eukaryotic translation elongation factor), apart from previously reported *CTNNB1, TP53* and *ALB* genes (26). However, given the high heterogeneity of HCC populations due to race, risk factors etc., a consensus list of driver genes among different HCC cohorts are yet to be identified. Moreover, the impact of driver mutations on HCC phenotypes, such as gene expression, have not been adequately investigated.

To address these issues, we have collectively conducted multi-modal meta-analysis on six HCC cohorts. The multi-modal data were collected from different approaches, ranging from WES/WGS data, RNA-Seq data, microRNA-Seq data to clinical data. We performed statistical analysis that combines the results of these cohorts, to derive 10 most significant consensus driver genes with significant functional impacts. To examine the association between driver mutations and gene expression, we built linear regression models using driver mutation and copy number variation (CNV) as predictors, and gene expression and miRNA (miR) expression as responses. Subsequent KEGG pathways and network analysis for these genes identified alterations in a broad spectrum of functions ranging from metabolic pathways, cell cycle to signaling pathways, as well as functional differences among the mutually exclusive driver genes. At the phenotypic level, we observed that consensus putative driver genes are predictive of survival differences among patients from cohorts with survival data. Some putative driver genes are significantly associated with physiological and clinical characteristics such as gender and age. In summary, we present the comprehensive picture of the functional relevance of driver genes in HCC, from molecular to phenotypic levels.

## Materials and Methods

### Dataset and processing

We used public domain HCC data from The Cancer Genome Atlas (TCGA) available at Genomic Data Commons (GDC) data portal, as of March 2017. In total, RNA-Seq, CNV and miR-Seq data comprise 371, 371 and 369 tumor samples, respectively. We used the R package TCGA-Assembler (v2.0) (27) to download the TCGA data. The mRNA-Seq data are represented as the normalized gene expression RSEM (RNA-Seq by Expectation Maximization) quantification values obtained from Illumina HiSeq assay platform, while miR-Seq data include ‘reads per million miR mapped’ (RPM) quantification values from Illumina HiSeq assay platform. CNV data represent gene-level copy number values obtained by taking the average copy number of genomic regions of a gene from the Affymetrix SNP Array 6.0 assay platform. To handle the missing values, we performed three steps. First, we removed the biological features (i.e. genes/miRs) if they were missing in more than 20% of the samples. Similarly, we removed the samples if they were missing for more than 20% of the features. Second, we used k-nearest neighbor based imputation using R *impute* package (28) to fill out the missing values. Last, we removed the genes with very low expression values (i.e. with RSEM/RPM< = 10 in the remaining samples). For TCGA mutation profile, the comprehensive Mutation Annotation File (LIHC-TP.final_analysis_set.maf) was downloaded from the FireBrowse portal of the Broad institute. We retrieved 362 samples (with HCC histology) having paired tumor and normal adjacent tissue WES data. Additionally, we obtained WES data from Liver Cancer (France): LICA-FR (n = 236), Liver Cancer (NCC, Japan): LINC-JP (n = 244) and Liver Cancer (China): LICA-CN (n = 163) cohorts, and WGS data from Liver Cancer (RIKEN, Japan): LIRI-JP (n = 258), all available as simple somatic mutation files from the International Cancer Genome Consortium (ICGC) web portal (29). These data from ICGC liver cohorts were published in the previous studies (16,22,30). Besides ICGC, we obtained another WES dataset (KOREAN (n = 231) from the early-stage HCCs (patients with surgical resection) with clinical information of patients published earlier (18).

### Consensus driver genes detection

To achieve the pool of consensus driver genes among six cohorts, we implemented the IntOGen platform (v3.0.6) (31), a comprehensive standalone pipeline for the identification of driver genes. The mutation profiles, from six cohorts, were subjected to MutSigCV (v1.4) (32) and OncodriveFM (33), both incorporated in the IntOGen pipeline. MutSigCV represents an advanced version of MutSig tool, which seeks to identify genes with significant positive selection during tumorigenesis. It calculates the personalized and gene-specific background random mutation rates, along with the implementation of expression levels and replication times as covariate factors. Complementarily, OncodriveFM uncovers the significant mutation space by applying the functional impact-based positive selection to identify the driver genes. We opted for a two-step screening to identify consensus drivers: a) we performed the q-value based screening, and followed by b) combined adjusted p-value based screening. For q-value based screening, we identified the genes from each module (i.e. MutSigCV and OncodriveFM) which satisfied: (i) q-values less than the threshold cut-off (q<0.1) in at least 3 of 6 cohorts, and (ii) mean q-value less than the threshold cut-off (q<0.1), across the cohorts. We obtained a set of “common drivers” by taking the intersection of the genes found in two modules. We chose the threshold of q-value<0.1 for both MutSigCV and OncodriveFM according to earlier studies (32,34). In order to make consensus drivers selection more stringent, we calculated the adjusted p-values for both MutSigCV-p-values and OncodriveFM-p-values for every cohort. For each of the “common drivers” identified in the previous step (q-value based screening) we conducted Fisher’s method for combined p-values and identified final “consensus driver genes” having significant combined p-values <0.05 for both MutSigCV and OncodriveFM. For downstream analyses, we excluded intergenic and intronic mutations.

### Determination of mutual exclusivity and co-occurrence

For each pair of consensus driver genes, we determined their association based on Fisher’s exact test with a p-value <0.05. For significant associations, if the log odds ratio was more than 0 for a pair of genes, the pair was called “co-occurred”, else “exclusive”. To detect the mutational exclusivity among gene sets (i.e. more than two genes), we applied the Dendrix algorithm (35) which is specialized to fish out gene sets with high coverage and exclusivity across the samples. We used gene set numbers k = 4, k = 5 and calculated their maximum weight with consideration of mutated genes and samples. We ran 100,000 iterations using Markov chain Monte Carlo approach to calculate empirical p-values for the top gene sets with the maximum weight.

For each cohort, we also used the bipartite graph to represent the mutations in the driver genes for each patient, using the patients and the driver genes as the distinct set of nodes. We used ForceAtlas2, a graph layout algorithm implemented in Gephi (36), to spatialize the graphs for mutual exclusivity. To compute the distances of the different cohorts the approach used was as follows: using the bipartite graph of each cohort, we computed the PageRank scores, a measure reflecting the connectivity of a node in a network (37), of the 10 driver genes. We used these scores as features representing cohorts. We then used Ward’s minimum variance method to cluster both the genes and the PageRank scores.

### Modeling relationships between consensus driver and gene expression

We made a binary (1, 0) matrix to indicate the mutation status of consensus driver genes in all samples. A value of 1 means the existence of at least one variant within the gene body, in the categories of nonsense, missense, inframe indel, frameshift, silent, splice site, transcription starting site and nonstop mutation. Otherwise, 0 was assigned to the gene. We made another table of CNV data similarly. We used voom function (*limma* package in R) to transform RSEM data prior to the linear modeling (38), then fit the linear models by minimizing generalized least squares similar to others (39). These linear models consider the effects of mutations of multiple consensus driver genes (predictors) and their CNVs on expression values of individual genes (responses) as follows:

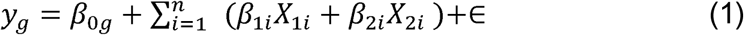

Where *y*_*g*_ is the vector representing expression value of gene *g* across all the *n* samples, *β*_0*g*_ is that baseline value of *g, X*_1*i*_ and *X*_2*i*_ are the mutation status and CNV of the consensus driver gene *i* (*i* = 1, 2….n), *β*_l_ and *β*_2_ are coefficients associated with the mutation status and CNV of the same gene, respectively. We performed multiple hypothesis tests on the significance values of the coefficients across all the genes using Benjamin – Hochberg (BH) adjustment, to determine the significant association between the driver genes and expression of all the genes (BH adjusted p-value <0.05). The accuracy of the applied tests and correction schemes was verified using a permutation approach, where each covariate was randomly permuted, breaking all correlations between genotype and expression. The permutation approach confirmed that the relationship between the pairs were significant, rather than being “random”.

### Pathway enrichment and network analysis

We conducted pathway enrichment analysis of the genes associated with somatic mutations and CNVs, using R package clusterProfiler (40). We used BH adjusted p-value = 0.05 as threshold to select the over-represented KEGG pathways. We used Gephi (36) based bipartite graphs to visualize driver gene-enriched pathways network.

### Modeling relationships between consensus drivers and miR expression

To find the relationship between driver genes (mutation and CNV) and miR expression, we implemented the linear model similar to that of equation (1). Here driver genes’ mutation and CNV status were treated as independent variables and miR expression as the response variable. To narrow down miRs that directly target these 10 drivers, we mined miRDB resource (41), which houses the miR-target interactions predicted by MirTarget (42) based on CLIP-Ligation experiments.

### Survival analysis of driver mutations

We used the Cox proportional hazards (Cox-PH) model (43) implemented in R *survival* package for the overall survival (OS) analysis of consensus driver genes. We developed Cox-PH model to fit the overall effect of all 10 driver genes on OS, with or without adjustments of clinical and physiological parameters (e.g. age, gender, grade etc.). For this, we used R *glmnet* package (44), since it enables penalization through ridge regression. We performed cross-validation to obtain the optimal regularization hyperparameter. The hyperparameter was selected by minimizing the mean cross-validated partial likelihood. To evaluate the performance of the survival models (45), we calculated the concordance index (CI) using function *concordance.index* in R *survcomp* package (46), based on Harrell’s C-statistics (47). We dichotomized the samples into high-and low-risk groups based on the median prognosis index (PI) score, the fitted survival values of the Cox-PH model (48 – 50). In the case of ties for the median PI, we shuffled the samples and randomly assigned them to either risk groups. We plotted the Kaplan-Meier survival curves for the two risk groups and calculated the log-rank p-value of the survival difference between them. We performed the similar survival analysis by adjusting the Cox-PH model with different physiological and clinical factors (e.g. age, gender, grade and tumor stage.

## Results

### Detection of consensus driver genes

To identify the consensus pool of driver genes among multiple cohorts of diverse populations, we used paired tumor-normal tissue of HCC WES data from TCGA as well as five other cohorts (WES/WGS). The clinical summary of patients in these 6 cohorts is provided (**Table S1)**. We assessed mutation significance and functional impact of protein coding genes using MutSigCV and OncodriveFM modules implemented in the IntOGen pipeline (see Materials and Methods) (**Figure 1A**). We identified the driver genes among the individual cohorts with the stringent threshold i.e. q-value <0.1 for both MutSigCV and OncodriveFM. Among these cohorts, TCGA contains the maximum number of drivers (20), while LICA-CN has 3 drivers only. LINC-JP, LIRI-JP, LICA-FR and KOREAN cohorts comprise 13, 11, 12 and 7 driver genes, respectively. *TP53* and *AXIN1* are two the driver genes shared by all the 6 cohorts (**Figure 1B**).

**Figure 1:**
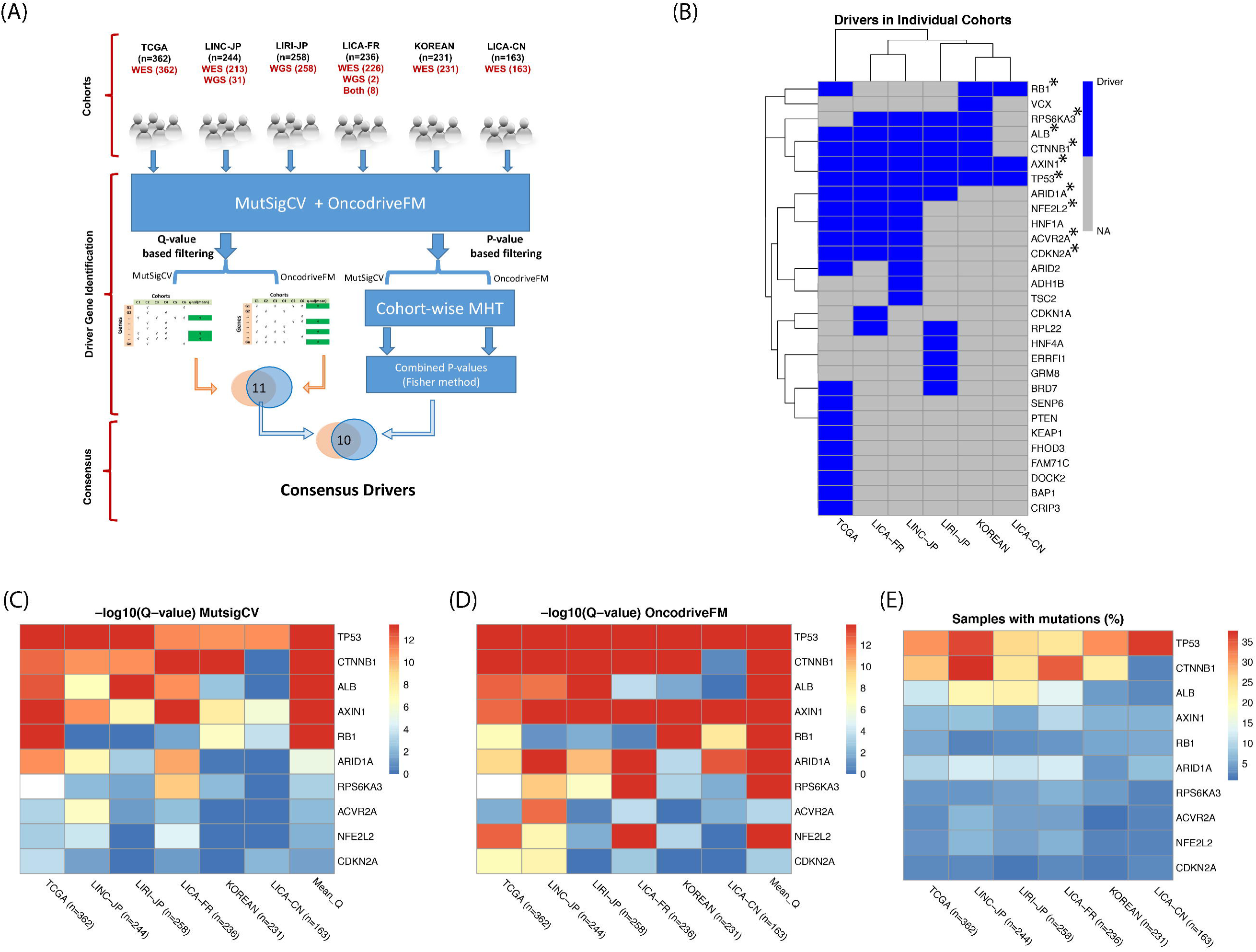
Consensus driver genes in 6 HCC cohorts. (A) IntOGen pipeline to identify consensus driver genes. (B) Driver genes from individual cohorts. Genes with asterisks represent consensus drivers. (C) Final 10 genes with mean q-value <0.1 from MutSigCV module. (D) Same 10 genes with mean q – value <0.1 from OncodriveFM module. (E) Percentage of sample coverage of driver gene mutations.

Next, we set out to define the “consensus driver gene”, which satisfied both of the following criteria: a) q-value based screening, where the mean q-value of a driver was less than the threshold cut-off (q<0.1) across the cohorts, and b) p-value based screening where Fisher’s combined adjusted p-value was less than 0.05 (**Figure 1A**). As a result, we identified 10 out of total 29 genes as “consensus driver genes” (**Figure 1B**). Interestingly, among patients with mutations in N consensus drivers (N = 0, 1, 2, 3, …, 10), single driver mutation (N = 1) is most frequently observed in all 5 cohorts, except LICA-CN cohort (**Figure S1**). Among these 10 genes, TP53 and CTNNB1 are most significantly mutated and functionally impactful genes based on q-values (**Figures 1C, 1D**), consistent with the earlier observations (18,21). However, some low-frequency mutation genes also have significant rankings per MutSigCV (**Figure 1C**). For examples, *CDKN2A, NFE2L2* and *ACVR2A* are all significant (mean q-values: 4.1e-02, 1.3e-02 and 6.1e-03 respectively), although their average mutation frequencies are less than 5% (**Figure 1E**). Thus, this workflow efficiently detects less frequent but consistently important driver genes.

### Analysis of consensus driver genes among cohorts

Next, we explored the mutation exclusivity status among these 10 driver genes across different populations (**Figure 2A**). As mentioned earlier, mutations from a single driver was most frequently observed in general (except LICA-CN). For patients with mutations in at least 2 consensus drivers, the fraction varies among cohorts: TCGA (26.5%), LINC-JP (42.6%), LIRI-JP (27.5%), LICA-FR (37.3%), KOREAN (16%) and LICA-CN (17.2%) (**Figure S1**). We used colored tiles in the plot to represent the specific type of mutation (e.g. missense, silent, frame shift etc.). A similar trend of mutation distribution exists in TCGA, three ICGC cohorts with large sample size (i.e. LINC-JP, LIRI-JP and LICA-FR) and the KOREAN cohort (**Figures 2A (i), (ii), (iii), (iv) and (v)**). Worth mentioning, LICA-CN cohort (n = 163) is most distinct from others and has the lowest *CTNNB1* mutation rate among all (**Figure 2A (vi)**). This exception may be attributable to HBV infection in LICA-CN cohort, as previous studies of HBV patients have reported the rare existence of *CTNNB1* mutations (22,23). In terms of the number of mutations per driver gene, most patients do not have too many (>25) mutations, except a very small fraction (**Figure S2**).

**Figure 2:**
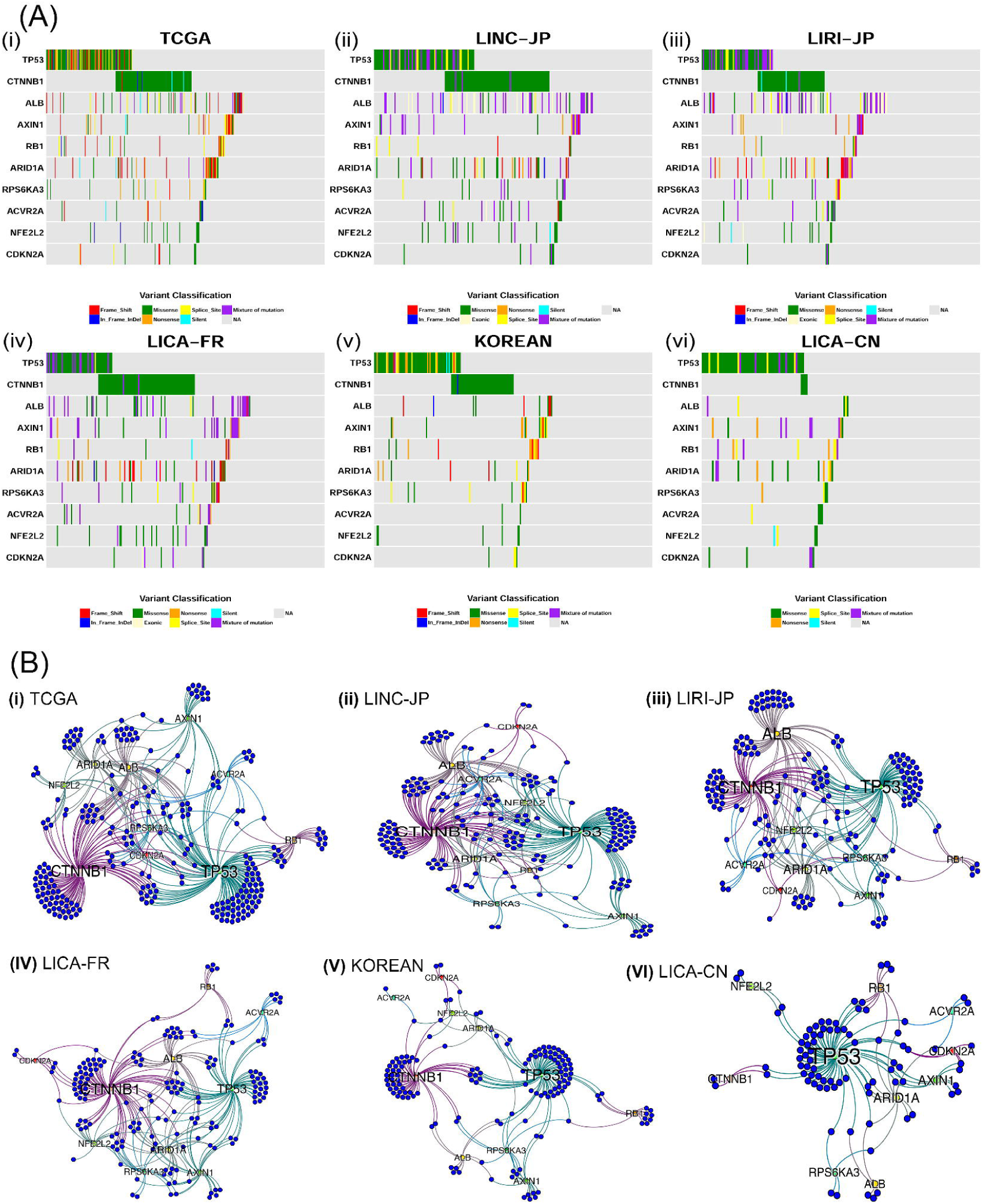
Mutual exclusivity among different driver genes in 6 HCC cohorts. (A) Co-mutation plots for the 6 HCC cohorts, where each colored tile represents one of the mutation types (i.e. frame shift, in-frame indel, missense, exonic, nonsense, splice site, silent or mixture of mutations) (i) TCGA (ii) LINC-JP (iii) LIRI-JP (iv) LICA-FR (v) KOREAN (vi) LICA-CN cohorts. (B) Bipartite graphs for mutual exclusivity of the same cohorts in (A). Blue nodes represent the patients and the other labeled nodes represent consensus driver genes, whose size is proportional to their degree.

Mutual exclusivity is apparent among some drivers (**Figure 2A**). For example, *CTNNB1* and *TP53* mutations are mutually exclusive in three of six cohorts, with significant Fisher’s exact test p-values in TCGA (P = 0.0303), LICA-FR (P = 0.0166) and KOREAN (P = 0.006). The mutual exclusivity between them was documented earlier (18). To detect mutual exclusivity beyond two genes, we used the Dendrix tool (35). Again, we observed significant mutational exclusivities (p-value = <0.05) for up to five genes in all 6 cohorts (**Figure S3**). *TP53, CTNNB1, RB1* and *AXIN1* and another cohort-specific genes are mutually exclusive in all five cohorts except LICA-CN. The other cohort-specific driver is *CDKN2A* (LINC-JP, LIRI-JP and KOREAN). Compared to the other five cohorts, LICA-CN cohort has most different five mutually exclusive drivers: *TP53, ACVR2A, ALB, CDKN2A*, and *RPS6KA3*.

We further visualized the relationships among patients, driver genes, and their topologies, using bipartite graphs (**Figure 2B**). The blue nodes and the labeled nodes represent patients and driver genes, respectively, and the edges between them indicate the existence of certain drivers in a particular patient. Based on the PageRank score that measures the connectivity and topologies of the graphs (see Materials and Methods), the similarity between TCGA and the other cohort descends in the following order: LINC-JP > LICA-FR > LIRI-JP > KOREAN > LICA-CN (**Figure S4**). KOREAN and LICA-CN cohorts are most distinct from other cohorts, with much fewer patients showing mutations in at least two driver genes. While KOREAN cohort mostly mutates in *TP53* and *CTNNB1* (however lacking *ALB* mutations like the other three cohorts), LICA-CN most dominantly mutates in *TP53* but not in *CTNNB1* or *ALB* (**Figures 2B (vi), S4**).

### The associations between gene expression and consensus driver gene mutation and CNV

To assess the associations between the genetics of consensus drivers and the transcriptome, we built generalized linear models using these driver genes’ mutation profile and their CNVs as the predictors, whereas gene expression values as the response variables, similar to other earlier genome-scale studies (51,52). These genetics based models decently predict gene expression values (R^2^ = 0.57) (**Figure 3A**), indicating that albeit the complex genetics and epigenetics regulatory mechanisms of gene expression, HCC driver gene mutations still convey important functional impacts on gene expression. Overall, our results show that around 62.5% (12,837) of genes are significantly associated (BH adjusted p-value <0.05) with these consensus driver genes. We list the number of genes significantly associated to each consensus driver gene in these linear models (**Figure 3B)**. The top two mutated genes are *CTNNB1* and *TP53* as expected, associated with over six thousand and nearly four thousand genes, respectively. Strikingly, the CNV of *ARID1A* is ranked 4^th^ and linked to expression changes in over 2,800 genes, despite its relatively low mutation rate of <10%.

**Figure 3:**
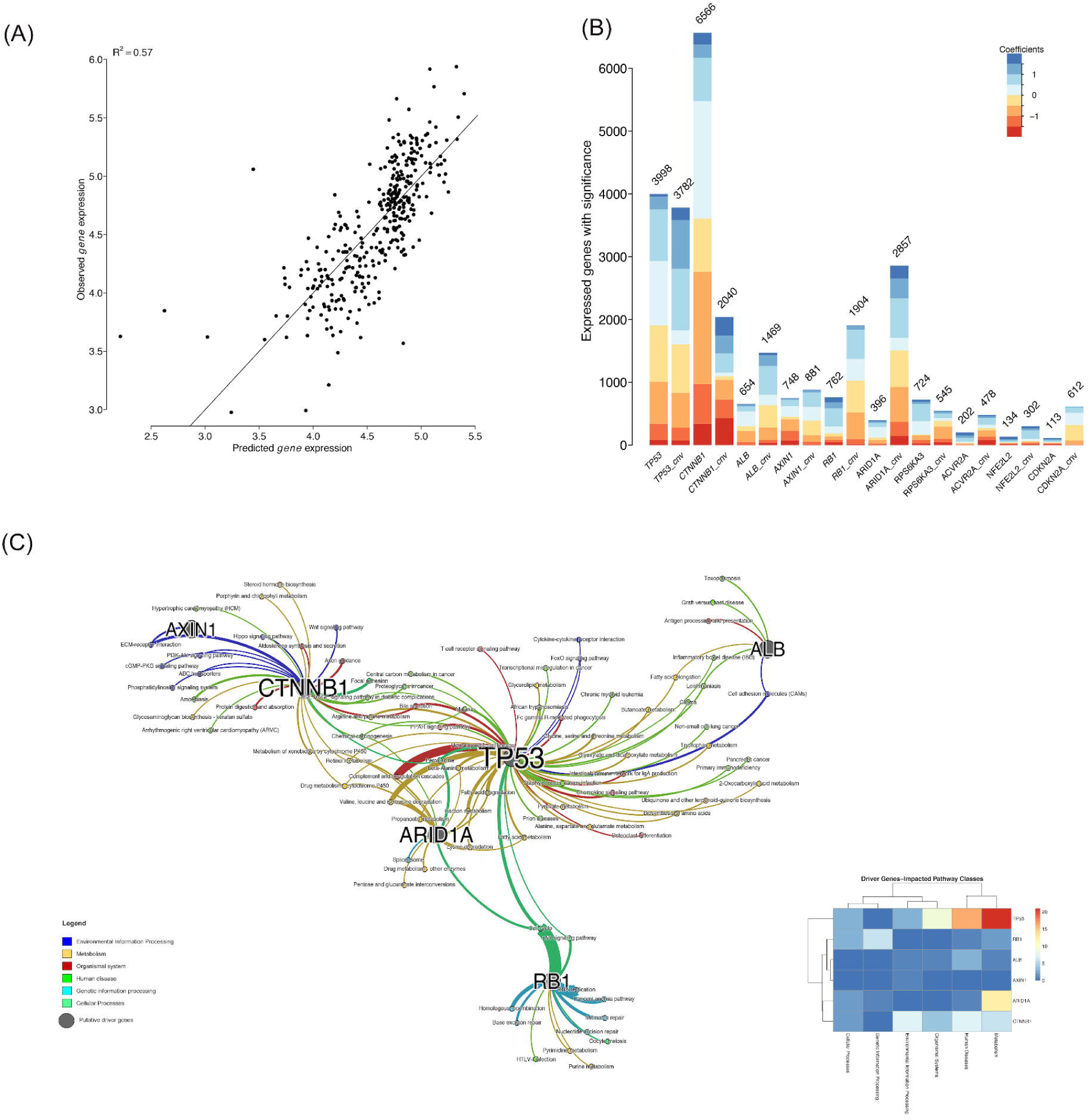
Associations of consensus driver genes with mRNA expression. (A) Correlation between observed and predicted gene expression. (B) The number of genes whose expression values are significantly associated with the driver gene mutation/CNV statuses. (C) Enriched KEGG pathways network among significant genes as shown in (B). The thickness of edges is proportional to the-log10 adjusted p-value.

To investigate the biological processes that these 12,837 genes are involved in, we conducted KEGG pathway enrichment analysis and detected 86 significantly (BH adjusted p-values <0.05) associated pathways (**Figure 3C**). We further categorized these pathways into 6 super groups according to the KEGG pathway organization, namely: cellular processes, environmental information processing, genetic information processing, metabolism, human diseases, and organismal systems (53). It is not surprising that the pathway super-group, affected most by the consensus driver genes, belongs to metabolic pathways. Among the driver genes, *TP53, CTNNB1*, and *ARID1A* are most densely connected to enriched pathways, due to the associations with gene expression changes. Some signaling pathways in the environmental information processing group are significantly influenced by driver genes, especially *CTNNB1*, which is associated with PI3K-Akt pathway, Wnt pathway and CGMP-PKG signaling pathway.

The association network between driver genes and pathways provide further support for mutual exclusivities observed earlier, at least partially, in that certain pathways are commonly associated by two mutually exclusive drivers. Between the well-known mutually exclusive *TP53* and *CTNNB1*, multiple pathways such as “bile secretion” and “proteoglycans in cancer” are shared. *TP53* and *ARID1A* are both involved in amino acid, carbon and fatty acid metabolism pathways. Heatmap of driver genes and six pathways classes (**Figure 3C**, insert) shows that *TP53* is associated with the maximum number of pathways related to Metabolism and Diseases, followed by *CTNNB1* and *ARID1A*.

We extended the linear modeling approach described earlier to examine the association between consensus driver genes and miRNA (miR) expression. Contrary to the vast prevalence of correlations between mRNAs and consensus drivers, we only found 167 miRs that are significantly associated with these drivers. Among them, 127 miRs are associated with driver gene CNV-level changes, 90 miRs are associated with the driver mutations, and 50 miRs are associated with both of them (**Figure S5**). This suggests that the major associations to protein coding genes are from driver mutations, not from non-coding regulatory elements miRs. The detailed association analysis between miR expression and consensus driver gene mutation/CNV is described in the supporting information (**File S1**).

### Associations between consensus driver genes and survival outcome

In order to test survival associations from all the driver mutations, we built multivariate Cox-PH models on overall survival in each of the four cohorts that have survival data (TCGA, LINC-JP, LIRI-JP and LICA-FR). We used the median prognostic index (PI) score generated from the Cox-PH model as the threshold (50), and divided samples into high and low risk groups accordingly (ties were assigned randomly to either risk group). The Kaplan-Meier survival curves of the two risk groups are presented for four cohorts (**Figure S6)**. For all the cohorts with survival data, the log-rank P-values between the Kaplan-Meier curves are significant (TCGA: P = 7e-03, C-index-0.58; LINC-JP: P = 5.3e-03, C-index = 0.67; LIRI-JP: P = 1.3e-02, C-index = 0.64 and LICA-FR: P = 3.4e-03, C-index = 0.61). To avoid potential confounding from age, gender, grade, stage in all 4 cohorts, we adjusted the Cox-PH model by these variables accordingly. Still, we identified significantly or almost significantly different survival groups (TCGA: P = 8e-03, LINC-JP: P = 2e-02, LIRI-JP: P = 7e-02 and LICA-FR: P = 4e-02) (**Figure S7**). All together, these results show that the driver genes’ mutational status is associated with HCC patients’ overall survival.

### Associations of consensus driver genes with health disparities

Previous studies have revealed aspects of disparities in HCC, such as preferable incidents in males (6,54). To reveal the possible link between these driver genes and gender/age, we conducted Fisher’s exact tests for gender, and Mann-Whitney-Wilcoxon tests for the continuous age variable. We found some significant associations of driver genes with gender and age (**Figure S8)**. To directly illustrate differences between categories, we calculated the relative risk (RR) of each category for the mutated genes vs. wild type genes. For age, RR was calculated after dichotomizing the samples based on mean age in the respective cohort.

With regard to gender, *CTNNB1*, a proto-oncogene, shows the most consistent evidence of preferred mutations in males, with an average RR = 1.2 in 5 cohorts (**Figures S8A-S8E**). Its strongest association comes from the TCGA cohort, based on significance level (P-value = 1.5e-05) and relative risk (RR = 1.4) (**Figure S8A**). Interestingly, *AXIN1* shows opposite and higher relative risks in females in 2 cohorts LIRI-JP (RR = 2.2) and KOREAN (RR = 2.2) (**Figures S8C, S8E**) and the overall average RR = 1.6 in 6 cohorts. Other drivers, such as *ALB* and *TP53*, are also preferred in males from 3 and 2 cohorts, respectively. For age, again *CTNNB1* is the driver gene with the strongest positive associations, for both relative risks (average RR = 1.2) and the number of cohorts (4 out of 6) (**Figures S8H-S8M**). Interestingly, *RB1* is the driver gene significantly and preferably prevalent in younger patients (3 out of 6 cohorts) (**Figures S8H, S8J and S8L**). However, *AXIN1* shows controversial associations with age between LINC-JP and LICA-FR cohorts, which may have to do with the different ethnicities between the two.

Additionally, TCGA and KOREAN cohort have information on risk factors of HCC (**Figures S8F, S8G**). The analysis shows that association between driver gene and risk factor is dependent on the cohort. In TCGA, *ACVR2A* shows significant higher RR among patients with fatty liver disease, alcohol users, and alcohol + HCV affected patients (**Figure S8F**). However, in KOREAN cohort, *CTNNB1* is the driver that shows significant associations in patients with HCV virus infection or no virus infection; however lower RR in patients with HBV infection (**Figure S8H**). Such difference in driver genes may be attributed to other factors, such as ethnicities or life styles. In terms of associations with race, *TP53* and *CDKN2A* both show higher RRs in Asians but lower RR in white (**Figure S8N**). For African Americans, RR for *TP53* is very high (RR = 2.2) however extremely low for *CDKN2A*.

## Discussion

In this study, we have pushed forward our understanding of the molecular and clinical associations of HCC drivers using multiple cohorts. Despite the heterogeneity among the datasets, we identified ten consensus driver genes derived from HCC WES/WGS data. Anchoring on these consensus driver genes, we investigated in-depth their transcriptomic and phenotypic associations, and prognostic values at the systems level. Detailed molecular mechanisms for each consensus drivers, although of interest to follow up, are not the focus of this multi-modal meta-analysis report.

A major contribution of this study is to associate the drivers with transcriptomic changes, which was previously unknown. The mutations and CNV of these consensus driver genes are correlated to around 63% mRNA transcriptome. These associated genes are involved in various pathways in cell cycle and DNA repair, metabolism, and signaling transduction. Interestingly, network analysis results show that mutually exclusively mutated genes have effects on some common biological processes, which may explain why mutations in both genes do not usually co-occur within the same patient. Surprisingly, only about 9% of miRs are associated with the consensus drivers globally, suggesting the major and direct role of driver mutations is on protein coding genes rather than regulatory components such as miRs. The survival plots based on the 10 consensus drivers’ mutation status alone show significant prognostic values, although not better than gene expression or protein expression as prognostic markers. In Pathology Atlas, the signatures are at the protein level, downstream of the phenotype (gene expression) that we consider here (55). The protein level change is the “output” reflective of many levels of regulations, from genetics, epigenetics, transcriptional and post-transcriptional modifications. Thus they are much closer prognostic biomarker for clinical phenotypes (such as survival) than driver mutations. For the purpose of optimizing prognostic biomarkers for HCC, we have recently reported another computational method based on deep learning (56) which takes the idea of integrating multi-omics datasets (57).

Our analysis reveals some unusual findings on genes with low mutation frequencies. One of them is that the CNV of *ARID1A* is one of the most “effective” events in the driver genes, prevalently associated with transcriptomic changes of 2,857 genes. *ARID1A* is a chromatin remodeller which is involved in transcriptional activation and considered as tumor suppressor (58). Previously, this gene is reported to be frequently deleted in HCC (20,59). *ARID1A*, a tumor suppressor gene, is depleted in advanced HCC and hence promotes angiogenesis via angiopoietin-2 (Ang2). *ARID1A*-deficient HCCs have been suggested as a good target for anti-angiogenesis therapies e.g. using sorafenib (60). *ARID1A* mutations in HCC have been reported to be associated HCC progression and metastasis in HBV-and alcohol-related HCC (23,61). Other infrequently mutated genes *such as ACVR2A* have also been reported in individual studies (16,30). Our stringent criteria for selection of consensus driver genes among 6 HCC cohorts highlights these low-mutated genes with consensus, reflecting that these may play a crucial role in HCC etiology. Along with *TP53* and *CTNNB1, ARID1A* stands out with densely connected sub networks.

Most interestingly, we have found evidence that some driver mutations are associated with gender and age disparities among HCC patients. *CTNNB1* is more prevalent in males, and it increases with age. Additionally, *TP53* and *ALB* are also more frequently mutated in males. Oppositely, *AXIN1* is more mutated in females. *AXIN1* encodes tumor suppressor gene axin-1, which is part of the beta-catenin destruction complex required for regulating *CTNNB1* levels through phosphorylation and ubiquitination (62). The opposite trend of gender association between *AXIN1* and *CTNNB1* can be explained by their antagonism relationship. Unexpectedly, we found that driver gene *RB1* is reversely related to ages of HCC patients. We do not know the etiology of such reversal age dependency of *RB1* in HCC. However, it has been well known that mutations in both alleles of the *RB1* gene are essential for retinoblastoma, which is often diagnosed in neonates (63). Additionally, drivers *TP53* and *CDKN2A* driver genes show such high RR in Asians but lower RR in White in TCGA data. However, the extrapolation of this observation as a general conclusion awaits for confirmations from more cohort studies having multiple races.

Unfortunately, the driver mutations in HCC are not yet designed as drug targets. Patients with advanced HCC have currently only two options of chemo-therapy: Sorafenib as the first line treatment and Regorafenib as the second line treatment (64). Both have very limited life-span extension, and both are multi-kinase inhibitors mainly targeting BRAF, which is not one of the consensus drivers we found. Therefore, we anticipate that this study presented here will motivate the development of therapeutic strategies that antagonize the most prevalent genetic mutations in HCC, by targeting genes including *TP53, CTNNB1* and *ARID1A*. In addition, other potential targets might be biological pathways that are enriched with driver mutations, such as Wnt/Beta-catenin pathway (with *CTNNB1*) and P53/cell-cycle pathway (with *TP53* and *RB1* drivers).

In summary, we have identified a consensus list of 10 driver genes in HCC, as well as their associations with downstream transcriptome and patient outcomes, such as survival, age and gender. Albeit the heterogeneity and complexity of HCC, the driver genes have broad and significant associations with global gene expression and molecular pathway functions, suggesting that HCC are genetically dominated diseases. Thus, this study provides an important and refined reference list for driver genes, which may serve as candidates for targeted therapies currently severely lacking in HCC.

## Availability of data and material

All HCC data are downloaded from the TCGA portal (https://tcga-data.nci.nih.gov/tcga/ and https://portal.gdc.cancer.gov/), FireBrowse portal from Broad institute (http://firebrowse.org/) and ICGC portal (https://dcc.icgc.org/).

## Acknowledgements

The authors would also like to thank Dr. Herbert Yu, Dr. Maarit Tiirikainen at University of Hawaii Cancer Center, Dr. Kenneth Kinzler at Johns Hopkins University, as well as all other group members in Garmire lab for helpful discussions and suggestions.

## Authors’ Contributions

LG conceived the project. KC and LL performed data analysis and wrote the manuscript, with the help of OP, SH and TC. All authors have read and revised the manuscript.

## Supporting Information

### Supplementary Figures

**Supplementary Figure S1: Distributions of the number of mutated consensus drivers in the six cohorts.**

**Supplementary Figure S2: Heatmaps of mutation counts in consensus drivers across samples in the six cohorts.** Red and blue colors represent low and high mutation counts in a particular driver.

**Supplementary Figure S3:** Bipartite graph showing the distribution of mutually exclusive genes identified with Dendrix across the 6 cohorts. Purple and yellow nodes represent genes and cohorts, respectively where the size of the node is proportional to the connectivity.

**Supplementary Figure S4: Similarity of the bipartite graphs among 6 HCC cohorts based on pageRank scores.** The darker color represents more predominant mutations.

**Supplementary Figure S5: Associations of consensus driver genes with miR expression.** Bipartite graphs representing (A) correlations between miR expression and mutations of the genes. (B) Correlations between miR expression and CNV. Green color represents positive correlation and red for anti-correlation.

**Supplementary Figure S6: Kaplan-Meier estimates of overall survival (OS) in 4 HCC cohorts.** A Cox-PH regression was used to build the overall survival model featuring the driver genes mutation profile. The samples were dichotomized into high and low risk groups by the median Prognostic Index (PI). (A) TCGA (B) LINC-JP (C) LIRI-JP (D) LICA-FR cohorts.

**Supplementary Figure S7: Kaplan-Meier of overall survival (OS) after adjustment for gender, age, stage and grade.** (A) TCGA (B) LINC-JP (C) LIRI-JP (D) LICA-FR cohorts.

**Supplementary Figure S8: Associations of gender and age with driver genes.** Shown are subsets of consensus driver genes with significant associations with gender (Fisher’s exact test with p-value<0.05) or age (Mann-Whitney-Wilcoxon test with p-value<0.05). Age is divided to two groups by the mean value (in parentheses) in each cohort.

### Supplementary Table

**Supplementary Table S1:** Clinical summary of 6 HCC cohorts.

### Supplementary File

**Supplementary File S1:** Associations between miR expression and consensus driver gene mutation/CNV.

